# Views and Experiences of Parents and Physicians on the Care Provided to Children with Sickle Cell Disease in Cameroon

**DOI:** 10.1101/2020.02.29.971218

**Authors:** Ettamba Agborndip, Benjamin Momo Kadia, Domin Sone Majunda Ekaney, Lawrence Tanyi Mbuagbaw, David C Rees, Helen Bedford

## Abstract

**Background:** Sickle Cell Disease (SCD) affects two in 100 Cameroonian new-borns, with 50-90% of affected children dying before their fifth birthday. Despite this burden, there is no national SCD programme in Cameroon. This study aimed to assess parents’ and physicians’ knowledge of SCD, their satisfaction with the quality of care and their recommendations to improve the treatment of SCD in Cameroon.

**Methods:** A multi-centre cross-sectional survey was conducted in English and French, using structured questionnaires distributed in electronic format to physicians throughout Cameroon. Paper-based questionnaires were also administered to parents in the West and North West regions of Cameroon. Data were entered into Microsoft Excel and analysed using the SPSS statistical software.

**Results:** Fifty-four parents and 205 physicians were recruited. We found that 72.2% of parents had good knowledge of SCD, 72.2% of parents were satisfied with the quality of care. Attending a sickle cell clinic (A*OR 22, 95% CI 17.70-250*) was significantly associated with having good knowledge. Just 14.2% of physicians had good knowledge and 23.3% of physicians were satisfied with the available management standards of SCD. Seeing more than five patients per month (*AOR 3.17, 95% CI 1.23-8.20*) was significantly associated with having good knowledge. Sickle cell clinics, national guidelines and subsidised treatment were the top three measures proposed by physicians and parents to improve the management of SCD.

**Conclusion:** Knowledge of SCD and satisfaction with care were poor among Cameroonian physicians. There is a need for a national programme and a comprehensive system of care for SCD in Cameroon.

## Introduction

Sickle cell disease (SCD) is the commonest monogenic disorder in the world. Sickle cell anaemia (SCA) is the most common variant of SCD in Africa and is lethal without treatment (1). In Africa, 50-90% children with SCA die before their fifth birthday, accounting for 5-16% of under-five mortality in some SSA countries (2).

In 2006, the World Health Organisation (WHO) declared SCD a global public health problem and recommended national control programmes and comprehensive health care management (CHCM) for SCD. This involves new-born screening (NBS), routine follow up, infection prevention, parental education and capacity building for health practitioners (3). While this has been widely implemented in high-income countries with good results, many SSA countries, including Cameroon, are still to introduce CHCM for SCD (4).

Cameroon is a central African country with about 22 million people. About 30% of the population is heterozygous for SCD, and 2-3% of children have SCD. The disease is severe in Cameroonian children, marked by recurrent crises and severe complications. According to Andong *et al* (5), children with sickle cell anaemia in Cameroon could experience up to seven painful episodes per year, and by the age of 14, 25% have either developed a stroke, chronic leg ulcers, nocturnal enuresis or gallstones. The severity of the disease in Cameroon has been linked to high temperatures, which favour dehydration and vaso-occlusive events; high malaria prevalence and the absence of CHCM (6).

Concerning health financing, Cameroon is yet to introduce universal health coverage. Patients depend on out-of-pocket spending, consuming more than 10% of their family income in most cases. In Cameroon, over 50% of rural and 30% of urban families are poor. Thus, most families are unable to keep up with the health demands of children with SCD. This results in high mortality rates, as SCD accounts for about 16% of under-five deaths in the country (6). Furthermore, there are disparities in health-seeking behaviour, with the poor and rural dwellers more likely to seek care from traditional doctors and drug peddlers (7). As a result, these children end up with more severe complications than those from affluent backgrounds.

Given the absence of a CHCM system in Cameroon, treatment of SCD mostly depends on the initiative of individual health providers. Comprehensive care for SCD is available in some private and missionary hospitals, in the form of monthly sickle cell clinics. However, the same cannot be said for most public hospitals, which serve the majority of Cameroon’s population. In the absence of a system to regulate the care of patients with SCD, management is largely dependent on personal knowledge and practice of the physicians. However, many physicians find it difficult to stay updated with current knowledge and practices, as SCD forms only a small portion of their clinical practice (8).

### Specific objectives

1. To assess parents’ and physicians’ knowledge of SCD
2. To determine the level of satisfaction of parents and physicians with the current standards of care for children with sickle cell anaemia in Cameroon
3. To identify the services recommended by parents and physicians to improve the quality of care for children with sickle cell anaemia in Cameroon.

## Participants and methods

### Operational terms

In this study, SCD refers to SCA.

### Study design

This study was a cross-sectional, self-administered (physicians)/ interviewer-administered (parents) questionnaire-based study done between June and August 2018.

### Study setting

Parents were recruited from the West and North West Regions of Cameroon, where the prevalence of SCD is said to be highest. Two health facilities were involved in the study, Fumbot District Hospital, a public hospital in the West with no sickle cell clinic, and the Association of People with SCD (ASPICAM) clinic in the North-West. ASPICAM clinic is coordinated by a medical doctor and three nurses and sees about 75 children with sickle cell anaemia per month. In this clinic, monthly meetings are organised for parents and SCD patients, during which are given health education, free folic acid and pain, medications, as well as counselling if necessary.

### Study procedure

#### Training

The principal investigator attended the certified Good Clinical Practice training as well as a seminar on New Sickle Cell Disease Trials in the UK.

#### Ethics

Ethical approval was obtained from the Institutional Review Boards of the Faculty of Health Sciences, University of Buea (Reference number: 2018/2238/UB/SG/IRB/FHS) and University College London (Reference number: 12751/001).

#### Study instrument

Two structured questionnaires were designed, one for physicians and the other for parents of children with sickle cell anaemia. The questionnaires contained similar sections, but the questions were slightly different for physicians and parents. The parents’ questions were formulated in simple language to ease their understanding. Physicians’ questionnaires were produced in English and translated into French, as these are the two official languages in Cameroon. The questionnaires were then pre-tested on a small sample of physicians. The questionnaires were anonymous and no identifying information was collected from participants.

#### Study population and sampling

using convenience sampling, participants were recruited consecutively until the end of the study period.

### Inclusion Criteria

1. Cameroonian Physicians who attend to children with sickle cell anaemia
2. Parents or carers of children aged 0-18 old with sickle cell disease
3. Children with sickle cell disease aged 13- 18 years old

### Exclusion criteria

1. Guardians who are not regular carers for children with sickle cell disease
2. Questionnaire with incomplete responses.

#### Recruitment

Due to socio-political instability, administrative authorities and the ethics committee discouraged the principal investigator (PI) from travelling to the study sites. Electronic versions of the questionnaires were sent via email to be printed and administered by two fifth-year medical students who had completed their paediatric rotations and had a good understanding of SCD and trained by the PI through skype.

Nurses identified parents during out-patient consultations, or sickle cell clinics. The medical students then introduced the study to them, presented and explained the information sheet. Consent was then obtained and they administered the questionnaire. At the end of each day, there was had a 30-minute Skype session, to assess completeness of questionnaires and discuss any issues. All questionnaires were emailed to the PI for data storage and analysis.

The physicians’ questionnaire was converted to an online survey to reach physicians in areas particularly affected by the ongoing political crisis. Filter questions were used to ensure that only participants who agreed to have understood the contents of the information sheet and gave consent were included in the study. The online survey was designed such that each participant could only respond to the survey once. Physicians’ contacts were obtained from the “Medical Forum” and the “Medical Syndicate”, to which majority of Cameroonian doctors belong. They were then contacted on a daily basis through personal emails and WhatsApp, with a short message inviting them to participate in the study and the link to the online survey.

### Statistical analysis

There were 10 questions on practice, 9 on knowledge, and 9 on satisfaction. Each question was given a score of 1. The sections on knowledge and practice comprised multiple choice and multiple (six) answer questions. For the multiple-choice questions, physicians and parents were given a score of one for each correct answer. For the multiple answer questions, parents were given a score of one if they ticked three or more correct answers, and physicians were given a score of one *only* if they ticked *all* the correct answers. In each section, the total score was categorised into good and poor as follows:

Knowledge: 0-5= poor; 6-9= good
Practice: 0-7=poor; 8-10= good
Satisfaction: 0-5=poor; 6-9=good

Data were analysed using SPSS version 22. Means and standard deviations were used to describe normally distributed variables, medians and interquartile ranges were used for non-normally distributed variables, and categorical variables were compared using the Chi-squared test. A p-value of < 0.05 was considered statistically significant. Logistic regression modelling was used to explore the effect of possible predictors of knowledge, practice and satisfaction. The frequencies and percentages of the measures recommended by physicians and parents to improve the management of SCD were calculated and represented on graphs.

## Results

### Baseline Characteristics of Respondents

A total of 205 physicians and 54 parents were recruited. Tables 1 and 2 show their characteristics.

### Characteristics of Physicians

Of the 205 physicians, 65.4% (135) were male, 70.2% (144) worked in a public hospital, and 84.9% (174) had been practicing medicine for under five years. The number of Children with SCD they reported seeing ranged between 0 and 100 per month, with a median of two children seen by each physician every month.

### Characteristics of Parents

Of the 54 parents (of 53 children), 87% (47) were female, 48.1% (25) had completed secondary education, and 77.4% (41) were unskilled workers. The mean age of parents was 42 years ± 12 years and the mean age of children was 11 years ± 5 years. About 90% of children attended school, each child had an average of two crises, had missed about seven days of school, and parents spent an average of 56,000 francs CFA (£77) on treatment every month (Table 2).

### Knowledge of SCD

Two hundred and five physicians responded to questions assessing knowledge, and 14.2% (29) had good knowledge of SCD. Most physicians knew the pathophysiology (84.4%) and the diagnostic test (99%) for SCD, 15.6% correctly identified all the triggers of sickle cell crisis, and about a third correctly identified the ideal age at which SCD should be diagnosed (Figure 1). There was strong evidence of an association between knowledge and the number of children seen. Physicians who saw more than five children with SCA per month were three times more likely to have good knowledge scores (AOR *3.17, 95% CI 1.28-8.20*) (Table 3).

Of the 54 parents who participated in this study, 72.2% (39) had good knowledge of SCD. Attending a sickle cell clinic (*p* = 0.005) and having secondary education (*p* = 0.026) were significantly associated with having good knowledge (Table 4). There was weak evidence for a difference in knowledge by gender (*p*=0.06), but this effect was lost after adjusting for other variables (logistic regression model). Parents who attended a sickle cell clinic were 22 times more likely to have good knowledge of SCD (*AOR 18.9, 95% CI 18.5-100*) than those who did not.

### Satisfaction with Care of Children with SCD

Sixty-two per cent of physicians were satisfied with the management of SCD in their health facilities. However, about two-thirds of physicians held the opinion that SCD is not properly managed in Cameroon. Most parents (72%) were satisfied with the standards of care provided to their children. It is worth noting, however, that despite this high rate of satisfaction among the parents 41 of the 54 (74.9%) of them were significantly financially set back by their children’s health expenses. Attending a sickle cell clinic appeared to be significantly associated with satisfaction. Parents who attended sickle cell clinics with their children were 19 times more likely to be satisfied with the care offered to their children that those who did not (*OR* 18.9, 95% CI 1.99 – 166).

### Recommendations for SCD Management

According to physicians, the top three measures needed to improve the management of SCD in Cameroon, were sickle cell clinics (67.8%), national guidelines (57.6%) and subsidised treatment (56.6%). Additionally, about 40% of physicians recommended NBS. For parents, the top three recommendations were setting up accessible sickle cell clinics (37.7%), provision of free (32.1%) and subsidised treatment (17%). A few parents (13.3%) recommended newborn screening.

## Discussion

### Summary of main findings

This study shows that the median age at which children with SCD were diagnosed was two years, they had about two crises and missed about seven days of school each month. Also, parents spent about 56,000 francs CFA (£77) every month on SCD-related costs. Knowledge of SCD among physicians was poor, with less than 15% having good knowledge. Physicians who saw more than five children with SCD per month were more likely to have better knowledge scores than those who saw fewer patients. However, knowledge of SCD was good among parents of Children with SCD, as 72.2% of them had good knowledge. Attending a sickle cell clinic and having secondary education were significantly associated with better knowledge. There was a significant difference in the organisation of care and physician practice, between the private and public sector. There were more services available for children with SCD, and practice scores were better among physicians in private hospitals than public hospitals. Most physicians (61%) were unsatisfied with the management of children with SCD in Cameroon. However, most parents were satisfied with the care their children received, but about 75% of them were uncomfortable with the monthly cost of care. Sickle cell clinics, national guidelines and subsidised treatment were the top three measures recommended by physicians to improve the management of SCD in Cameroon. The main recommendations made by parents were setting up sickle cell clinics and provision of free/ subsidised treatment.

### Knowledge of SCD

Physicians had poor knowledge of SCD. Most of them failed to identify the correct age at which SCD should be diagnosed, and only 15% of them correctly identified all the triggers of crises. Early diagnosis and linkage to care, as well as prevention of crises, are key aspects in the management of SCD. If physicians do not know the triggers of SCD crises, they cannot teach parents how to avoid them. This results in recurrent crises, reflected in this study by the high frequency of crises and number of school days missed reported by parents.

Poor knowledge among physicians could also explain the high frequency of acute and chronic complications among Cameroonian children with sickle cell anaemia, as reported by Andong, Ngouadjeu (5), who found a high prevalence of stroke, leg ulcers and refractive eye diseases among children with SCD in the Littoral and South West regions of Cameroon. The low level of knowledge among physicians highlights a need for training or CME on SCD.

The problem of poor knowledge of SCD among physicians is not limited to Cameroon. Dennis-Antwi, Dyson (9) and Gomes, Vieira (10) also reported poor knowledge of SCD among physicians in Ghana and Brazil respectively. Given that SCD is endemic in these countries, it is expected that physicians should have a good grasp of the disease and its management. This is however not the case. This poor knowledge among physicians could be attributed to the fact that, despite the high prevalence in these countries, SCD forms only a small portion of their practice (as seen in this study, where 84.5% of physicians saw less than 5 children with SCD per month). As such, physicians are more likely to develop expertise in more common diseases. This is demonstrated in our study by the fact that doctors who saw more than five children with SCD per month had significantly better knowledge scores, with similar findings reported by Vieira (10) and Tanner (11) among family physicians in Brazil and USA, respectively.

Up to 72% of parents had good knowledge of SCD. This is in contrast to the findings of Hassan, Awwalu (12) in Nigeria, who reported that most (60%) parents had poor knowledge of SCD. This difference could be because the majority of the parents in our study attended a sickle cell clinic, where they received regular and up-to-date information about SCD, while the parents in the Nigerian study attended primary care facilities with no sickle cell clinics. The good knowledge among parents in our study is similar the findings of by Rahimy, Gangbo (13), among parents attending a sickle cell clinic in Benin, thus highlighting the advantage of a comprehensive system of care for children with SCD. Families in our study may not be representative of SCA families in Cameroon as a whole in that they were already known to SCA clinics. It is likely that overall knowledge of SCA amongst patients and parents in Cameroon is much lower.

### Satisfaction with Care

Our study demonstrated a significant difference in the organisation of care of children with SCD in public and private health facilities. Private health facilities seemed to pay more attention to these children, as demonstrated by scheduled visits (planned care), the presence of registries for children with SCD and the more frequent use of guidelines, which according to participants were mostly local guidelines. This difference was reflected in physicians’ practice, as private physicians were more likely to recommend NBS and prescribe correct routine medications and vaccines than public physicians.

Overall, however, very few physicians routinely prescribed penicillin V (40/174) and malaria prophylaxis (18/174) for children with SCD. These findings differ from those reported by Galadanci, Wudil (14) in Nigeria, where malaria prophylaxis was routinely prescribed in all, and penicillin V in 72% of health facilities. Considering that malaria is a common trigger of sickle cell crises and an important cause of mortality among these children these findings are disturbing and highlight the absence of a system to regulate the management of SCD in Cameroon.

In this study, about two-thirds of physicians were dissatisfied with standards of care for SCD in Cameroon. Some of the reasons we identified include the absence of a national programme and guidelines for the management of SCD in Cameroon, unavailability of basic diagnostic tests for SCD in several health facilities and the lack of psychosocial and financial support available for children with SCD and their families in Cameroon.

On the other hand, most parents (72%) were satisfied with the quality of care provided to their children. The findings of this study are similar to those reported by Israel-Aina, Odunvbun (15), who found high levels of satisfaction among parents in Benin city, Nigeria. They, however, differ from those of Hassan and colleagues (2017), who assessed the level of satisfaction among parents attending a primary health care centre in Nigeria. They reported low levels of satisfaction, with only 46% of parents rating the services as either good or excellent. This difference in findings could be because the parents in this study and the one by Israel-Aina *et al*., attended a sickle cell clinic, where they received regular psychosocial support as well as information on how to manage their children at home, whereas those in Hassan’s study received no such care.

Despite the high levels of satisfaction among parents in our study, it is worth noting that three-quarters of parents were uncomfortable with the monthly expenses on their children’s health. This could be the reason why subsidised treatment was one of the top recommendations made by parents. These parents also represented a pre-selected group who were already receiving some sort of specialist SCA care.

### Recommendations for SCD Management

The top three measures proposed by physicians to improve the management of SCD were sickle cell clinics, national guidelines and subsidised treatment. Similarly, parents proposed sickle cell clinics, free treatment and subsidised treatment.

The recommendation of sickle cell clinics is in accordance with the WHO recommendation of comprehensive care, especially in countries like Cameroon with a high prevalence of SCD (3). It also highlights the need for a system of care that caters to the specific needs of children with SCD, and the benefits of such a system cannot be overstated. Rahimy and colleagues (13) in Benin reported fewer crises and a 78% improvement in the physical and nutritional status of these patients following the initiation of comprehensive care for these in Benin.

The proposal of national guidelines by physicians in this study is similar to findings reported by Mainous, Tanner (11), who carried out a national survey to assess the attitudes of family physicians towards the management of SCD. In their study, about 70% of physicians indicated a need for clinical decision tools (guidelines). Treatment guidelines are an important tool for management of diseases, especially in paediatrics. Strict use and adherence to guidelines have been beneficial in reducing the morbidity and mortality of diseases like malaria (16), as well as reducing the cost for patients and their families by preventing complications (17). National guidelines would ensure good practice in the care and treatment of SCD.

Parents spent an average of 56,000 francs CFA (£77) per month, which is almost double the minimum wage (36,000 francs CFA) in Cameroon (18). With such financial constraints, parents may be unable to access essential care for their children, which contributes to frequent morbidity and a vicious cycle. The recommendation of providing free or subsidised treatment by parents and physicians draws attention to a possible significant lack of financial protection of children with SCD and their care givers. This points to an important gap in universal health in coverage in Cameroon.

### Strengths and limitations of this study

- Although sickle cell disease is very common in Cameroon (and Africa), very little is known about the perspectives of patients and healthcare providers regarding standards of SCD management in this setting. Previous studies generally provide a one-sided view, as they focused on either the parents or physicians. Our study provides more insight into gaps in SCD management by exploring the views and experiences of both users (parents/patients) and suppliers (physicians) of available SCD treatment services. Such balanced view of key stakeholders’ perspectives is crucial in the formulation of adequate national policies governing SCD management.
- This study does not only highlight the need for sickle cell clinics but goes further to demonstrate their effectiveness as shown by the better knowledge and satisfaction scores among parents who attended sickle cell clinics. This would potentially save policymakers the cost of carrying out feasibility studies and provide a model for national sickle cell clinics in Cameroon.
- CHCM recommended by the WHO has not been implemented in Cameroon, and the current standards of care are not commensurate with the burden of SCD. This study provides evidence on gaps in SCD management in the country. This will, in turn, inform the decisions of policy makers as they allocate resources and plan the introduction of a national comprehensive care programme for SCD management.
- Physicians were recruited via an online survey, which was more feasible considering time and cost constraints, coupled with the ongoing political instability at the time of the study. However, this approach introduced the possibility of selection bias, as physicians in remote areas with no internet access, who might have different views and experiences on the management of SCD from those in urban or semi-urban areas, were not sampled.
- The study did not distinguish between views of paediatricians and general practitioners. Nonetheless, the management of SCD in Cameroon is mainly by general practitioners, as there are few paediatricians in Cameroon. Thus, the views expressed by the participants in this study are likely to reflect the views of most physicians who manage children with SCD in the country

## Conclusion

The findings of this study highlight the importance of comprehensive care for children with SCD and the effectiveness of this strategy is evidenced by the high levels of knowledge and satisfaction among parents who attended sickle cell clinics. This study also demonstrates a need for national management guidelines, as well a system to regulate SCD management, as more knowledge did not translate into better practice among physicians. A national registry for SCD describing patterns of complications, disease burden and the natural history of SCD in Cameroon could be useful in monitoring and evaluating the national SCD program. NBS and subsidised treatment could be important in optimising the effectiveness of a national SCD program which we advocate for.

## Declarations

## List of Abbreviations

ACS: Acute Chest Syndrome
CCC: Comprehensive Clinical Care
CME: Continuing Medical Education
SCA: Sickle Cell Anaemia
DRC: Democratic Republic of Congo
FDA: Food and Drug Administration
Hb: Haemoglobin
HbS: Haemoglobin S
HLY: Healthy Life Years
LMIC: Low and Middle-income Countries
NBS: Newborn Screening
NO: Nitric Oxide
PI: Principal Investigator
RBC: Red Blood Cell
SCA: Sickle Cell Anaemia
SCD: Sickle Cell Disease
SSA: Sub-Saharan Africa
UK: United Kingdom
US: United States
VOC: Vaso-occlusive Crisis
WHO: World Health Organisation

## Authors’ contributions

EA: conception of the study, literature search, data management and statistical analysis. BMK: data collection and manuscript preparation and editing. DSME: reviewed and edited the study proposal and manuscript. LTM: critical review of the manuscript and provision of intellectual guidance. DCR: critically reviewed the study protocol and the initial manuscript for technical and intellectual consistency. HB: Conception of the study, critical review of the final manuscript, provided technical and intellectual guidance.

## Availability of data and material

Data can be made available by the corresponding author upon reasonable request.

## Acknowledgements

We are grateful to the following persons: Dr Nsah Bernard; Dr Njefi Kevin; Dr Ngassa Stewart; Dr Christian Akem Dimala; Dr Christie Linonge; Dr Yong Oryn; Mr Bekou Fokou Hygin; and Ms. Arrey Echi. Their input was invaluable.

## Compliance with Ethical Standards

### Ethical approval

All procedures performed in studies involving human participants were in accordance with the ethical standards of the institutional and/or national research committee and with the 1964 Helsinki declaration and its later amendments or comparable ethical standards.

### Informed consent

Informed consent was obtained from all individual participants included in the study

## List of Figures

Figure 1: Recruitment of study participants

Figure 1: Proportion of Physicians with Good Knowledge of Various Aspects of SCD

## List of Tables

Table 1: Characteristics of Physicians

Table 2: Characteristics of Parents and Their Children

Table 3: Predictors of Physicians’ Knowledge of SCD

Table 4: Predictors of Parents’ Knowledge

## References

1. Rees DC, Williams TN, Gladwin MT. Sickle-cell disease. The Lancet. 2010;376(9757):2018–31.

2. Grosse SD, Odame I, Atrash HK, Amendah DD, Piel FB, Williams TN. Sickle Cell Disease in Africa: A Neglected Cause of Early Childhood Mortality. American Journal of Preventive Medicine. 2011;41(6, Supplement 4):S398–S405.

3. WHO. Sickle-cell anaemia: Report by Secretariat. WORLD HEALTH ORGANIZATION 2006.

4. Noubouossie D, C. TT, B. C, P. NB, N. MD. Sickle-cell disease in sub-Saharan Africa. ISBT Science Series. 2016;11(S1):256–62.

5. Andong AM, Ngouadjeu EDT, Bekolo CE, Verla VS, Nebongo D, Mboue-Djieka Y, et al. Chronic complications and quality of life of patients living with sickle cell disease and receiving care in three hospitals in Cameroon: a cross-sectional study. BMC Hematology. 2017;17:7.

6. Wonkam A, Mnika K, Ngo Bitoungui VJ, Chetcha Chemegni B, Chimusa ER, Dandara C, et al. Clinical and genetic factors are associated with pain and hospitalisation rates in sickle cell anaemia in Cameroon. Br J Haematol. 2018;180(1):134–46.

7. Makoge V, Maat H, Vaandrager L, Koelen M. Health-Seeking Behaviour towards Poverty-Related Disease (PRDs): A Qualitative Study of People Living in Camps and on Campuses in Cameroon. PLoS neglected tropical diseases. 2017;11(1):e0005218.

8. Adewoyin AS. Management of Sickle Cell Disease: A Review for Physician Education in Nigeria (Sub-Saharan Africa). Anemia. 2015;2015:791498.

9. Dennis-Antwi JA, Dyson S, Ohene-Frempong K. Healthcare provision for sickle cell disease in Ghana: challenges for the African context. Diversity & Equality in Health and Care. 2008;5.

10. Gomes LMX, Vieira MM, Reis TC, Barbosa TLA, Caldeira AP. Knowledge of family health program practitioners in Brazil about sickle cell disease: a descriptive, cross-sectional study. BMC Family Practice. 2011;12:89.

11. Mainous AG, 3rd, Tanner RJ, Harle CA, Baker R, Shokar NK, Hulihan MM. Attitudes toward Management of Sickle Cell Disease and Its Complications: A National Survey of Academic Family Physicians. Anemia. 2015;2015:853835.

12. Hassan A, Awwalu S, Okpetu L, Waziri A. Knowledge and perception of patients with sickle-cell disease about primary care providers in Zaria, North-West Nigeria. Archives of Medicine and Surgery. 2017;2(1):12–5.

13. Rahimy MC, Gangbo A, Ahouignan G, Adjou R, Deguenon C, Goussanou S, et al. Effect of a comprehensive clinical care program on disease course in severely ill children with sickle cell anemia in a sub-Saharan African setting. Blood. 2003;102(3):834.

14. Galadanci N, Wudil BJ, Balogun TM, Ogunrinde GO, Akinsulie A, Hasan-Hanga F, et al. Current sickle cell disease management practices in Nigeria. International health. 2014;6(1):23–8.

15. Israel-Aina YT, Odunvbun ME, Aina-Israel O. Parental Satisfaction with Quality of Health Care of Children with Sickle Cell Disease at the University of Benin Teaching Hospital, Benin City. Journal of Community Medicine and Primary Health Care. 2017;29(2):33–45.

16. Biai S, Rodrigues A, Gomes M, Ribeiro I, Sodemann M, Alves F, et al. Reduced in-hospital mortality after improved management of children under 5 years admitted to hospital with malaria: randomised trial. BMJ (Clinical research ed). 2007;335(7625):862.

17. Tieder JS, Robertson A, Garrison MM. Pediatric hospital adherence to the standard of care for acute gastroenteritis. Pediatrics. 2009;124(6):e1081–7.

18. Foumena J-C. Minimum wage in Cameroon: PART I 2016 [Available from: https://www.linkedin.com/pulse/minimum-wage-cameroon-part-i-jean-claude-foumena.

