## Supplementary figures and images for "Views and Experiences of Parents and Physicians on the Care Provided to Children with Sickle Cell Disease in Cameroon"

### Supplemental figure 1

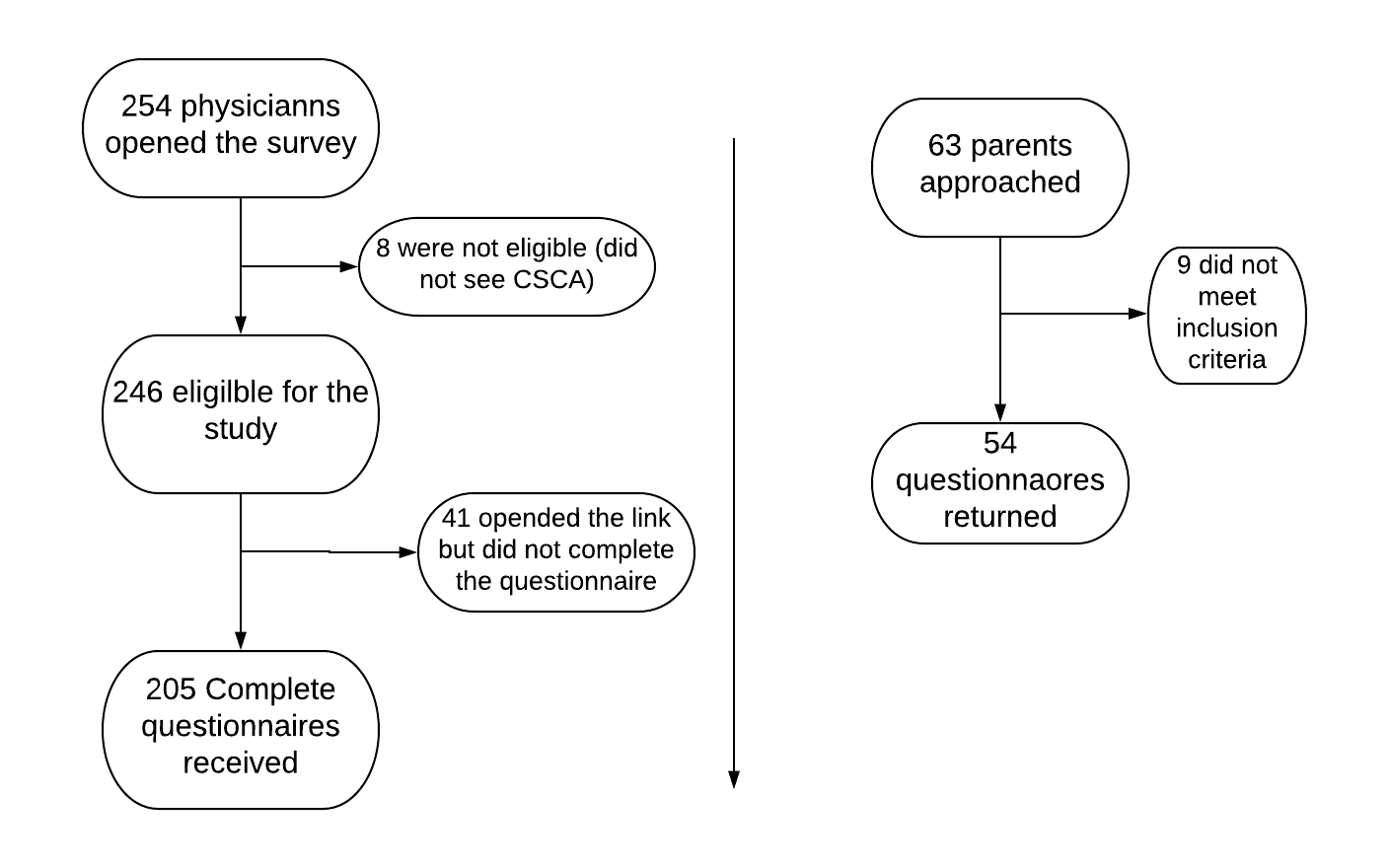

**Figure 1: Recruitment of study participants**
