## Supplemental figure 2 for "Views and Experiences of Parents and Physicians on the Care Provided to Children with Sickle Cell Disease in Cameroon"

N=205

Frequeny (%)

|  | Triggers | Ideal diagnostic age | Tranfusion Criteria | Intitial Symptom | Diagnostic test | Routine Drugs | Pathophysiology | Dental Visits | Ophthamology visits |
| --- | --- | --- | --- | --- | --- | --- | --- | --- | --- |
| 1 | 15.6 | 35.6 | 53.7 | 50.2 | 99 | 40.2 | 83.4 | 45.9 | 76.6 |

**Figure 2: Proportion of Physicians with Good Knowledge of Various Aspects of SCD**
