## Supplemental table 1 for "Views and Experiences of Parents and Physicians on the Care Provided to Children with Sickle Cell Disease in Cameroon"

**Table 1: Characteristics of Physicians**

| Numeric variables | | | |
| --- | --- | --- | --- |
|  | **Median** | | **Interquartile Range** |
| Age (years) | 29 | | 27-31 |
| Duration in Medical Practice (years) | 3 | | 2-5 |
| Number of Children with SCD seen per month | 2 | | 1-5 |
| Categorical Variables | | | |
|  | **Number (n)** | **Percentage (%)** | |
| Gender |  |  | |
| Female | 71 | 34.6 | |
| Male | 134 | 65.4 | |
| Type of Health facility |  |  | |
| Public | 144 | 70.2 | |
| Private | 61 | 29.8 | |
| Duration in Medical Practice |  |  | |
| < 5 years | 168 | 82 | |
| >5 years | 37 | 18 | |
| Number of Children with SCD seen per month |  |  | |
| < 5 | 174 | 84.9 | |
| >5 | 31 | 15.4 | |
| Region |  |  | |
| North West or South West | 102 | 49.8 | |
| Others | 103 | 50.2 | |
