## Supplemental table 2 for "Views and Experiences of Parents and Physicians on the Care Provided to Children with Sickle Cell Disease in Cameroon"

**Table 2: Characteristics of Parents and Their Children**

| Children | | |
| --- | --- | --- |
| Variable | **Median** | **Interquartile range** |
| Child’s age at diagnosis (years) | 1.8 | 0.7-6 |
|  | **Mean** | **Standard Deviation** |
| Child’s age (years) | 11 | 5.40 |
| Number of school days missed per month | 7 | 6.5 |
| Number of crises per month | 2 | 1.96 |
| Child attends school | **Frequency** | **Percentage** |
| Yes | 48 | 90.6 |
| No | 5 | 9.4 |
| Parents | | |
|  | **Mean** | **Standard Deviation** |
| Carer’s age (years) | 41.8 | 12.34 |
| Monthly expenditure on SCD (Francs CFA) | 56156 (‎£77) | 40000 (‎£55) |
|  | **Frequency** | **Percentage** |
| Carer’s gender |  |  |
| Female | 47 | 87.0 |
| Male | 7 | 13.0 |
| Type of occupation |  |  |
| Skilled | 12 | 22.6 |
| Unskilled | 41 | 77.4 |
| Attends a sickle cell clinic |  |  |
| Yes | 46 | 88.9 |
| No | 6 | 11.1 |
| Level of education |  |  |
| None | 4 | 7.7 |
| Primary | 19 | 36.5 |
| Secondary | 25 | 48.1 |
| Tertiary | 4 | 07.4 |
