## Supplemental table 3 for "Views and Experiences of Parents and Physicians on the Care Provided to Children with Sickle Cell Disease in Cameroon"

**Table 3: Predictors of Physicians' Knowledge of SCD**

|  | | Univariable Predictors | | | | Multivariable Predictors | |
| --- | --- | --- | --- | --- | --- | --- | --- |
| Variable | | **Knowledge (n/%)** | | **p Value** | **OR (CI)** | **p Value** | **OR (CI)** |
|  |  | Poor | Good |  |  |  |  |
| Gender | |  |  |  |  |  |  |
|  | Male | 117 (87.3) | 17 (12.7) | 0.410 | 1.4 (0.627-3.123) |  |  |
|  | Female | 59 (83.1) | 12 (16.9) |  | 1 |  |  |
| Duration in medical practice | |  |  |  |  |  |  |
|  | <Five years | 144 (84.7) | 24 (14.3) | 0.903 | 1.067 |  |  |
|  | >Five years | 32 (85.6) | 5 (13.5) |  | 1 |  |  |
| Type of Health Facility | |  |  |  |  |  |  |
|  | Public | 124 (86.1) | 20 (13.9) | 0.871 | 1 |  |  |
|  | Private | 52 (85.2) | 9 (14.8) |  | 1.073 (0.458-2.512) |  |  |
| Number of Children with SCD seen | |  |  |  |  |  |  |
|  | < Five | 154 (85.5) | 20 (11.5) | **0.010**** | 1 | **0.017**** | **1** |
|  | >Five | 22 (71.0) | 9 (29.0) |  | 3.15 (1.28-7.81) |  | **3.17( 1.28-8.20)** |
