## Supplemental table 4 for "Views and Experiences of Parents and Physicians on the Care Provided to Children with Sickle Cell Disease in Cameroon"

**Table 4: Predictors of Parents' Knowledge**

|  | |  | Univariate Analysis | | | Multivariate Analysis | |
| --- | --- | --- | --- | --- | --- | --- | --- |
| Variable | | **Good (n/%)** | **Poor (n/%)** | **OR (95%CI)** | **P value** | **OR** | **P value** |
| Gender | |  |  |  |  |  |  |
|  | Female | 36 (76.6) | 11(23.4) | 4.36 (0.85-22.55) | 0.06 |  |  |
|  | Male | 03 (42.9) | 04 (51.7) | 1 |  |  |  |
| Relationship with child | |  |  |  |  |  |  |
|  | Mother | 29(78.4) | 08(21.6) | 1 | 0.14 |  |  |
|  | Other | 10(58.8) | 07(41.2) | 0.39(0.11-1.37) |  |  |  |
| Duration with the diagnosis | |  |  |  |  |  |  |
|  | <Five years | 12 (70.7) | 05 (29.4) | 0.89 (0.249-3.17) | 0.86 |  |  |
|  | >Five years | 27 (73.0) | 10 (27.0) |  |  |  |  |
| Attends Sickle Cell Clinic | |  |  |  |  |  |  |
|  | Yes | 38 (79.2) | 10 (20.8) | **18.9 (18.5-100)** | **0.005**** | **22 (17.70-250)** | **0.016**** |
|  | No | 1 (16.7) | 5 (83.3) | **1** |  | **1** |  |
| Level of Education | |  |  |  |  |  |  |
|  | None | 4 (100) | 0 (0) | **0.089** |  |  |  |
|  | Primary | 9 (47.4) | 10 (52.6) | **0.146** | **0.026**** | **0.147 (0.031-0.702)** | **0.016**** |
|  | Secondary | 22 (84.6) | 4 (15.4) | **1** |  | **1** |  |
|  | Tertiary | 3 (75.0) | 1 (25.0) | **0.545** |  |  |  |
